# Salicylic acid accumulation correlates with low anthocyanin production in Arabidopsis

**DOI:** 10.1101/2025.06.08.658514

**Authors:** Matěj Drs, Oksana Iakovenko, Jhonny Stalyn Orozco Hernandez, Pavla Beáta Trhlínová, Vedrana Marković, Viktor Žárský, Tamara Pečenková, Martin Janda

**Affiliations:** Department of Experimental Plant Biology, Faculty of Science, University of South Bohemia in České Budějovice, Branišovská 1645/31a, České Budějovice, 370 05, Czech Republic; Laboratory of Cell Biology, Institute of Experimental Botany, Czech Academy of Sciences, Rozvojová 263, Lysolaje, 165 02 Praha 6, Czech Republic; Department of Experimental Plant Biology, Faculty of Science, Charles University, Viničná 5, 128 44, Prague 2, Czech Republic

**Keywords:** AVI bodies, anthocyanins, salicylic acid mutants, *NahG*, exocyst, autophagy

## Abstract

Anthocyanins, flavonoid pigments, are essential photoprotective agents and play a pivotal role in enhancing plant resilience to environmental stressors. It has been shown that anthocyanin production is inhibited when pattern-triggered immunity (PTI) is activated in *Arabidopsis thaliana*. An important component of PTI is the phytohormone salicylic acid (SA). Interestingly, exogenous treatment with SA has been shown to induce anthocyanin content in grape, apple, maize roots, rose callus, or *Arabidopsis* seedlings. In this study, we used several *A. thaliana* mutants with modulated SA content to decipher the role of endogenous SA in anthocyanin production in *A. thaliana*. We treated WT and mutants with anthocyanin-inducible conditions and measured anthocyanin content using spectroscopy. We showed that high endogenous SA accumulation correlates with low anthocyanin production. This was confirmed by the treatment of the *A. thaliana* seedlings with exogenous SA. Additionally, using microscopy in the *5gt* mutant, which exhibits enhanced production of anthocyanin vesicular inclusions (AVIs) due to the inhibition of ligandin-dependent vacuolar import, we showed that high endogenous SA also correlates with lower AVI abundance. Comparative analysis of *Arabidopsis* WT and mutants used in this study indicates a possible inhibitory effect of SA accumulation on anthocyanin content under anthocyanin-inducible conditions (AICs). We suggest that under AICs, SA downstream signaling independent of NPR1 is responsible for lower anthocyanin accumulation.

**Highlights:**

- High endogenous SA correlates with low anthocyanin content under AIC in Arabidopsis
- SA signaling, not biosynthesis, is responsible for the inhibition of anthocyanins
- High SA concentration decreases the abundance of AVI bodies

**GRAPHICAL ABSTRACT:** 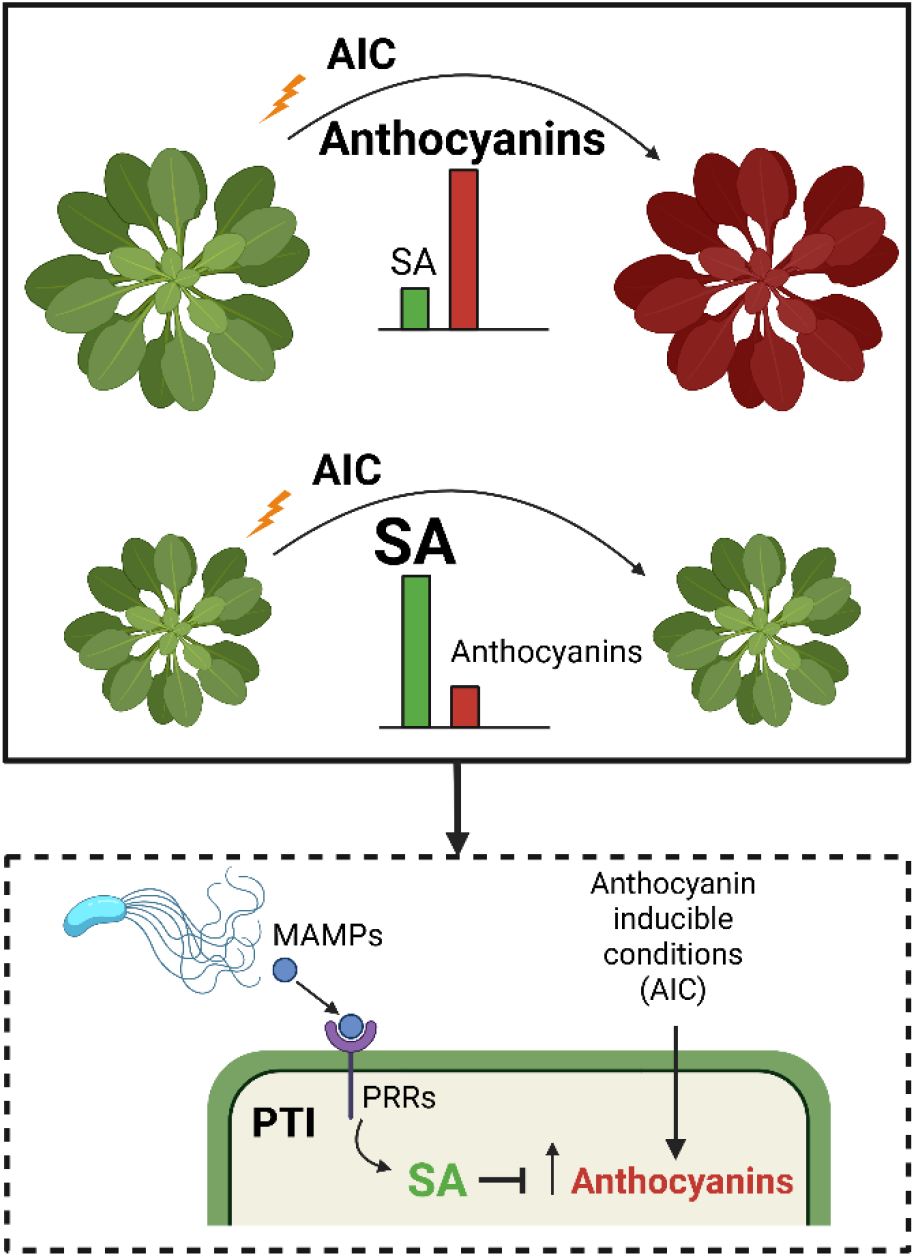

**Graphical abstract**. Anthocyanin-inducible conditions (AICs), such as changes in day length, high sucrose in the medium, or treatment with kinetin, trigger the biosynthesis of anthocyanins. Previously, it was shown that activated pattern-triggered immunity (PTI), caused by the recognition of microbe-associated molecular patterns (MAMPs) by pattern recognition receptors (PRRs), inhibits anthocyanin production. A typical PTI response is an increased production of salicylic acid (SA). In this study, we show that a high concentration of SA and its downstream signaling, rather than SA biosynthesis itself, reduces anthocyanin accumulation.

## 1 INTRODUCTION

Plants respond to various abiotic stresses, such as osmotic stress, UV light, and changes in day length, by inducing the synthesis of anthocyanins, which are water-soluble flavonoid pigments responsible for the characteristic red, blue, and purple colors of plant tissues. These pigments are synthesized via the phenylpropanoid pathway and represent glycosylated derivatives of anthocyanidins. Anthocyanidins are glycosylated at the 3-O and/or 5-O positions by the corresponding UDP-glucose:flavonoid 3-O-glucosyltransferase (3GT) or 5-O-glucosyltransferase (5GT), respectively. Once synthesized, anthocyanins accumulate in the large central vacuole, where their oxidation is prevented. Their synthesis occurs at the cytoplasmic surface of the endoplasmic reticulum (ER), from where they are transported to the vacuole. Over the past years, several models have been proposed to explain anthocyanin transport to the vacuole(Li and Ahammed, 2023).

The ligandin model suggests that anthocyanins are actively transported across the tonoplast by specific membrane importers, with the assistance of glutathione S-transferase (GST) enzymes (Poustka et al., 2007; Sun et al., 2012). The vesicular model proposes that anthocyanins, once imported into the ER lumen, are transported to the vacuole via vesicles (Gomez et al., 2011; Pourcel et al., 2010). Interestingly, the vesicular movement of anthocyanins—or anthocyanin vesicular inclusions (AVIs)— may also be regulated by microautophagy (Chanoca et al., 2015; Pourcel et al., 2010).

Anthocyanins have antioxidant properties, protecting plants against oxidative damage and high temperatures. Due to their coloration, anthocyanins also play an important role in attracting pollinators (Li and Ahammed, 2023). However, they are also induced by pathogen attack (Liu et al., 2020) and contribute to plant defense mechanisms by deterring herbivores and exhibiting antimicrobial properties (Lev-Yadun and Gould, 2008). On the other hand, it has been shown in *Arabidopsis thaliana* that activated pattern-triggered immunity (PTI), induced by treatment with the peptides elf18 and flg22, classical examples of microbe-associated molecular patterns (MAMPs), inhibits anthocyanin production triggered by high sucrose or UV-B irradiation (SCHENKE et al., 2011; Serrano et al., 2012; Zhou et al., 2017). Mitogen-Activated Protein Kinase Kinase 6 (MKK6) plays an important role in this inhibition (Wersch et al., 2018). A crucial integral component of PTI defence is also autophagy (REF), and autophagy loss-of-function mutants accumulate high levels of salicylic acid, exhibit decreased levels of anthocyanins, and show enhanced spontaneous hypersensitive reaction (HR) (Masclaux-Daubresse et al., 2014).

An integral component of PTI signaling is the phytohormone salicylic acid (SA) and its signaling pathway. SA concentration increases upon treatment with elf18 or flg22 (Tsuda et al., 2008). SA biosynthesis in plants occurs via two pathways, both starting from chorismate in the chloroplasts. The first is the isochorismate synthase (ICS)-dependent pathway, which is responsible for the majority of induced SA in *A. thaliana* during bacterial attack (Wildermuth et al., 2001). The second pathway depends on phenylalanine ammonia-lyase (PAL) (Dempsey et al., 2011). The ICS-dependent pathway appears to be more important under stress conditions in *Brassicaceae*, whereas in other plant species, the PAL pathway plays a more prominent role (Ullah et al., 2023). The PAL-dependent biosynthetic pathway shares its initial step with the flavonoid biosynthesis pathway, and multiple mutations in PAL genes result in decreased levels of both SA and anthocyanins (Zhang et al., 2021).

In this study, we focused on the role of the SA pathway in anthocyanin accumulation under abiotic stress conditions in *Arabidopsis thaliana*.

## 2 MATERIALS AND METHODS

### 2.1 Plant material

We used *Arabidopsis thaliana* ecotype Columbia col-0 as a wild type (WT), additionally we used *A. thaliana* mutants with Col-0 genetic background which we used in previous studies: *bon1-1, pi4kβ1/pi4kβ2, fah1/fah2, exo70B1-1, NahG, bon1-1/snc1-11, NahG/pi4kβ1/pi4kβ2* (Pluhařová et al., 2019), *tn2* (Zhao et al, 2015), *tn2/exo70B1-1* (Zhao et al, 2015); previously published, but not by us *5gt* (Chanoca et al., 2015) and which we generated in this study: *5gt/exo70B1-1* and *5gt/exo70B1-1/tn2*. Plants used for analyses were grown in three distinct growing conditions. Either *A. thaliana* plants were grown in hydroponia using ¼ Hoagland solution, and after 4-5 weeks, treated by anthocyanin-inducing conditions (AICs; particularly treated either with sucrose or kinetin). The temperature was set to 20 ± 2 °C, light intensity was 130 - 150 μmol.m^−2^.s^−1^, and photoperiod was 10 h / 14 h (light/dark) (these growing conditions were used for results in Fig. 1). Alternatively, plants were cultivated in Jiffy palettes for 5 weeks under the short day conditions (8/14; light intensity 100 μmol.m^−2^.s^−1^; 70% humidity); additionally, the seedlings were cultivated in liquid ½ MS in 6-wells plates (Deltalab, Spain) for 5 days under the long days conditions (16/8; light intensity 100 μmol.m^−2^.s^−1^).

**Figure 1.**
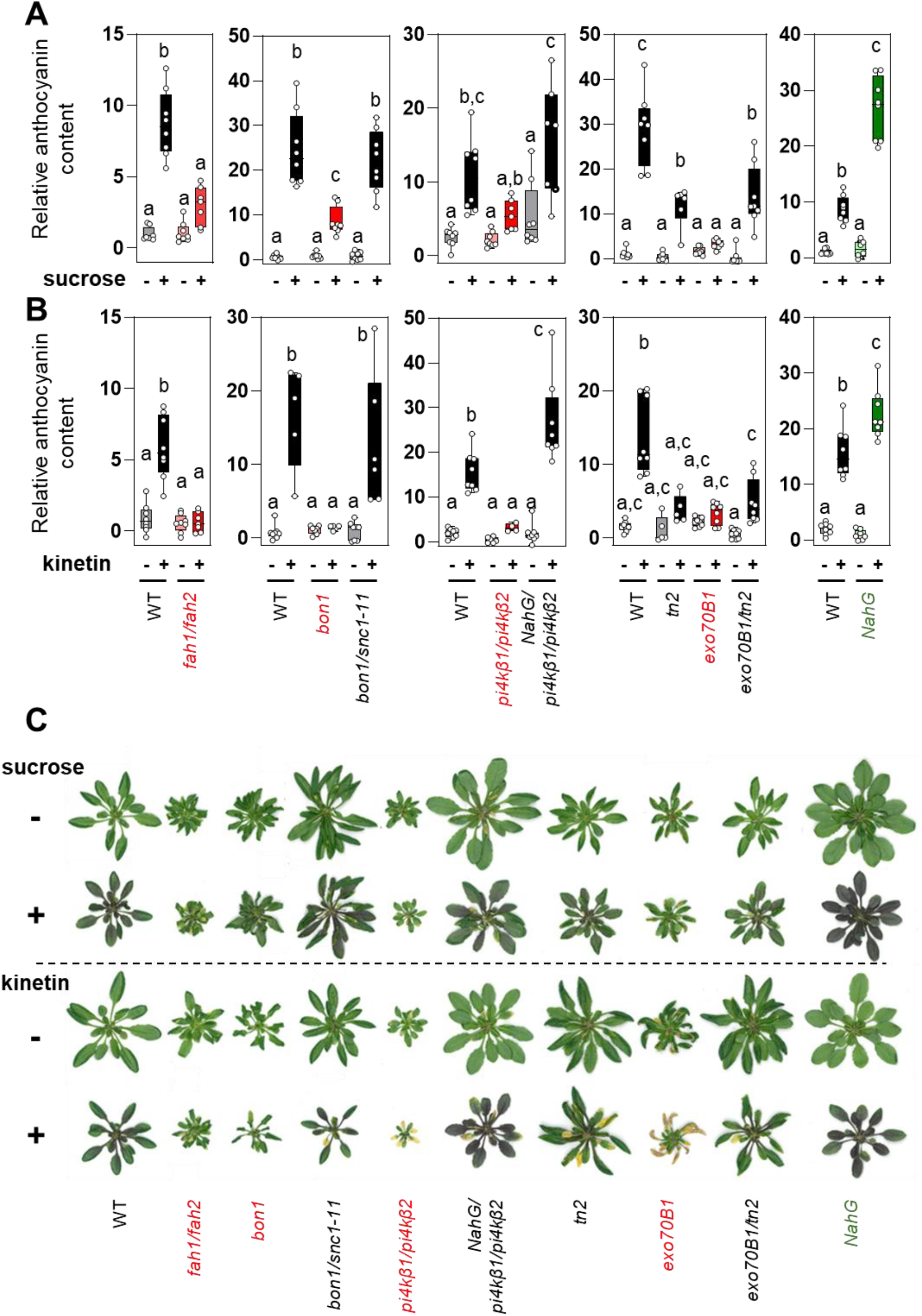
High endogenous salicylic acid inhibits anthocyanin production. Roots of 4-5-week-old *Arabidopsis* plants grown hydroponically were treated with **(A)** 7% sucrose for 72 hours and **(B)** 50 µM kinetin for 168 hours. **C)** Representative images of *Arabidopsis* plants after sucrose and kinetin treatment. Statistical analysis was performed using one-way ANOVA followed by a post hoc Tukey test (*p* < 0.05, *n* = 5–8 independent samples). Experiments were repeated 2–3 times for each genotype and treatment.

### 2.2 Anthocyanin-inducing conditions

For plants grown in hydroponia, the AIC 7% sucrose or 50 μM kinetin (diluted in 5M KOH) was added to fresh fresh-prepared ¼ Hoagland solution. A new ¼ Hoagland solution was used as a mock. For the kinetin experiment, the mock was ¼ Hoagland solution with KOH (50μl/L 5M KOH).

The induction of anthocyanins accumulation for Jiffy tablets cultivated plants was performed by the switch from the short day condition were transferred into a chamber with a long day (16/8; light intensity 100 μmol.m^−2^.s^−1^; 70% humidity) and cultivated for 14 days. The accumulation of anthocyanins in seedlings was triggered by adding of fresh or ½ MS with 7 % sucrose into cultivation wells with seedlings for 48 hours treatment (these growing conditions were used for the results in the Fig. 2 and 3).

**Figure 2.**
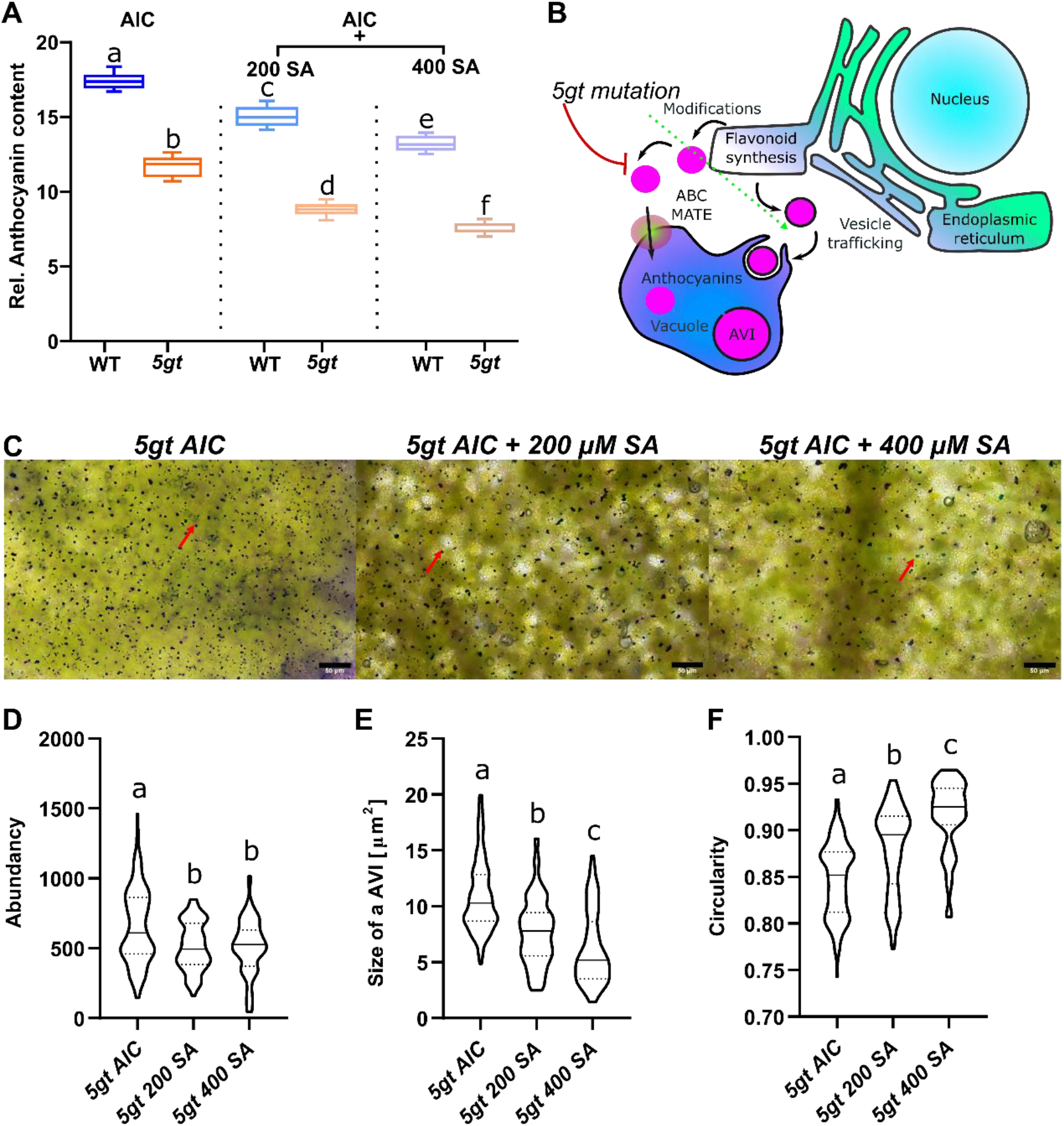
Exogenous SA inhibits anthocyanin production and alters vacuolar inclusion bodies. **A)** Plot showing the analysis of anthocyanin content in seedlings of WT and *5gt* mutant under anthocyanin-inducing conditions (AIC; 7% sucrose), combined with salicylic acid (SA) treatment at two concentrations (200 and 400 μM). **B)** Diagram illustrating the effect of the *5gt* mutation on anthocyanin transport. **C)** RGB images of cotyledon leaves under the same treatments as in **(A)**; scale bar = 50 μm (. Red arrows indicate AVIs (Anthocyanin Vacuolar Inclusions); Analysis of AVI properties: abundance **(D)**, size **(E)**, and circularity **(F)**. Statistical analysis was performed using one-way ANOVA followed by a post hoc Tukey test (*p* < 0.001). *n* = at least 16 independent leaves per genotype **(D, E, F)**; 12 seedlings per genotype per treatment for **(A)**.

**Figure 3.**
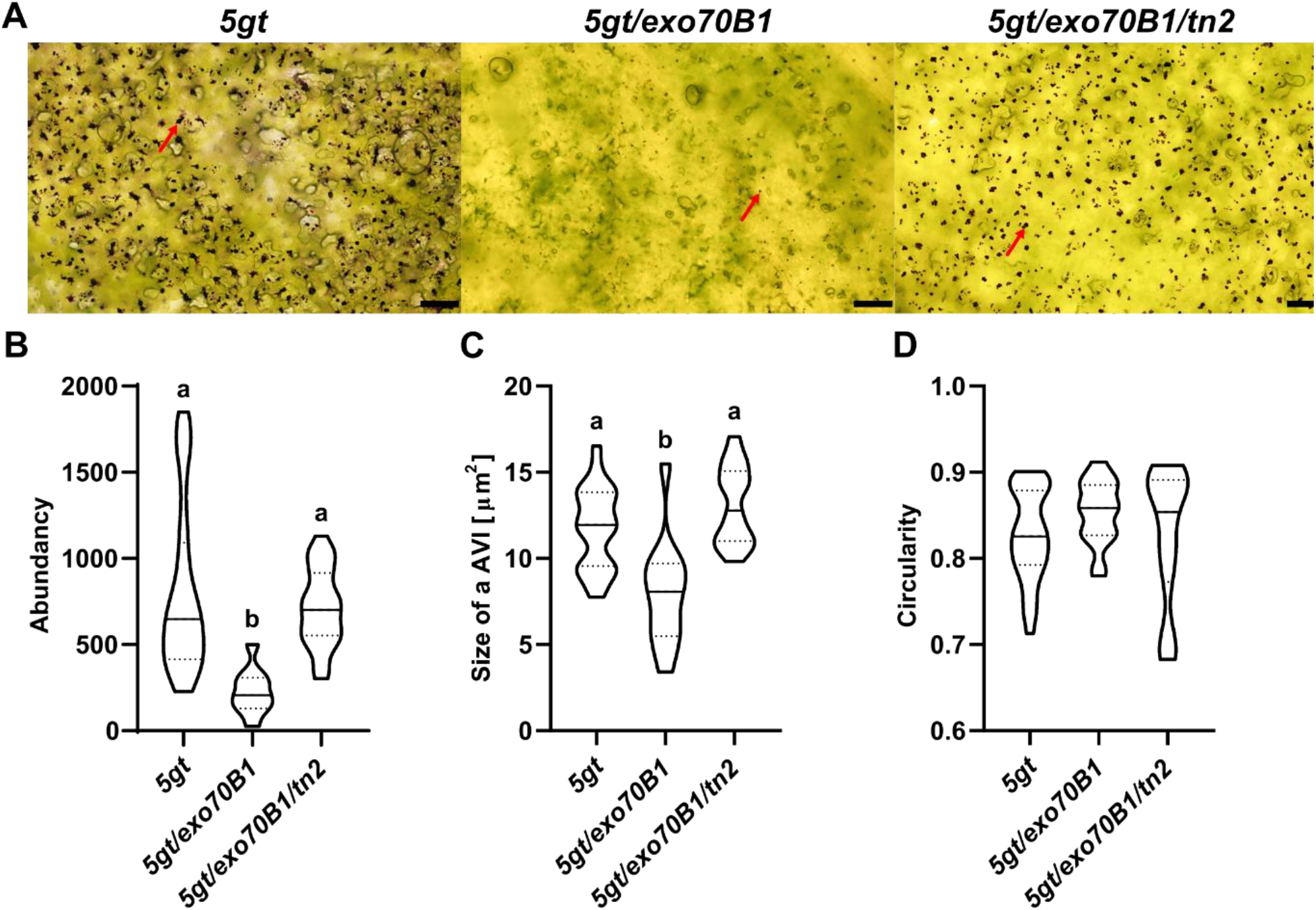
Endogenous SA inhibits anthocyanin production and alters vacuolar inclusion bodies. **A)** RGB images of adult leaves from 7-week-old plants following a short-day to long-day transition. Red arrows highlight AVIs (Anthocyanin Vacuolar Inclusions); scale bar = 50 μm. Quantitative analysis of AVI properties: abundance (**B**), size (**C**), and circularity (**D**). Statistical analysis was performed using ANOVA followed by Tukey’s post hoc test (p < 0.001, *n* = at least 16 independent leaves per genotype).

### 2.3 Salicylic acid treatment

At IEB 5-day-old seedlings grown in liquid ½ MS were treated in cultivation wells with ½ MS containing 7% sucrose with or without different concentrations of SA (either 200 or 400 µM) for 48 hours.

### 2.4 Anthocyanin analysis

#### 2.4.1 Spectrofotometry

For anthocyanins measurement from *A. thaliana* grown in hydroponia fully developed adult leaves were used. Detached leaves were homogenized in the extraction solution (methanol with 1% HCl), vortexed and centrifuged at 20,000 RCF for 20 min. Absorbance was measured using a spectrophotometer (Analytik Jena Specord 210 Plus), and anthocyanin content was calculated by the formula ((λ_528-_ λ_657_)/4)/FW (Zou et al., 2017). Anthocyanin isolation from whole seedlings was done with the same method.

#### 2.4.2 Microscopy

Cotyledons or mature leaves were mounted onto microscopic slides and observed using the Olympus (BX-51 with Olympus DP-74 camera) microscope with a real-time z-stack stitching plugin (Olympus-cellSens software). AVIs were analyzed using FIJI (Schindelin et al., 2012) with a labkit plugin to segment AVIs from the background.

## 3 RESULTS AND DISCUSSION

### 3.1 High endogenous salicylic acid reduces anthocyanin accumulation

Since previous reports on anthocyanin accumulation, conducted on various plant species and tissues under biotic stress conditions, have yielded conflicting results, we aimed to analyze the effect of both endogenous and exogenous SA on anthocyanin accumulation in *Arabidopsis thaliana* using distinct experimental approaches.

We treated plants cultivated in hydroponia with 7% sucrose or with 50μM kinetin, typical anthocyanin-inducing conditions (AICs) (**Fig. 1**). Under established AICs we tested collection of Arabidopsis mutants with distinct SA content changes: *fah1/fah2*; *bon1-1*; *pi4kβ1/pi4kβ2* and *exo70B1* with high SA content (Pluhařová et al., 2019) and corresponding double mutants *bon1-1/snc1-11*; *NahG/pi4kβ1/pikβ2* and *exo70B1/tn2* with normalized low SA content (Pluhařová et al., 2019; Zhao et al., 2015); for comparison we used WT and *NahG* with lowered SA content due to SA degradation (Šašek et al., 2014). *NahG* is a mutant expressing bacterial salicylate hydroxylase metabolizing SA to katechol (Delaney et al., 1994). Our results demonstrated that high endogenous SA correlates with low anthocyanin accumulation in AICs (**Fig. 1**). In all double mutants with normalized WT SA level (*bon1-1/snc1-11, exo70B1/tn2* and *NahG/pi4kβ11/pi4kβ2*), we observed restored anthocyanin production (**Fig. 1**). This reinforces the phenomenon that SA and anthocyanin accumulation negatively correlate (Masclaux-Daubresse et al., 2014). Results showing strong production of anthocyanins in *NahG/pi4kβ1/pikβ2* and certainly in *NahG* alone (**Fig. 1**), demonstrate that not SA biosynthesis flux but SA accumulation or SA downstream signalling is responsible for lower anthocyanin accumulation.

The master regulator of SA signalling is NPR1 protein (Janda and Ruelland, 2015). Kulich et al. (2013) showed that crossing of *npr1* knock-out mutant with *exo70B1* mutant did not restore anthocyanin accumulation in *exo70B1* under AICs (Kulich et al., 2013). Additionally, Zhao et al. (2015) showed that double mutant *npr1/exo70B1* has even higher content of SA than *exo70B1* single mutant (Zhao et al., 2015). Taken together, we suggest that SA alone is not an inhibitor of anthocyanin production, but SA signaling (e.g. changes in metabolic flux caused by SA), including NPR1-independent pathway, is responsible for the inhibition of anthocyanin accumulation under AICs. However, the detailed mechanism of that inhibition remains to be elucidated.

### 3.2 Exogenous application of SA decreases vacuolar anthocyanin accumulation, including anthocyanin vacuolar inclusion bodies (AVIs)

Mutant plants from our variable SA content collection provided information about the inhibitory effect of accumulation of endogenous SA on the ability to produce and accumulate anthocyanins (**Fig. 1**). However, it was shown that exogenous treatment with SA triggers increase of anthocyanin content in Arabidopsis (Liu et al., 2020), grape berries and cells (Khalili et al., 2022; OBINATA et al., 2003; Oraei et al., 2019; Yue et al., 2023), in pomegranate (García-Pastor et al., 2020), in pepper plants (Mahdavian et al., 2008), in *Rosa hybrida* calli (Ram et al., 2013), in maize roots (Jain and Srivastava, 1984), in combination with elevated CO_2_ also in ginger (Ghasemzadeh et al., 2012), in combination with sucrose in *Pistacia chinensis* leaves (Song et al., 2020), and SA alleviated the inhibitory effect of ethylene on anthocyanin production in *Brassica napus* (Tirani et al., 2013). Thus, we also externally applied different concentrations of SA together with sucrose on seedlings and we observed surprisingly a direct inhibitory effect of exogenous SA on the amount of anthocyanins under AIC in concentration-dependent manner in *A. thaliana* seedlings (**Fig. 2A**). Along with the spectroscopy we used also a second approach for the quantification of anthocyanins vacuolar accumulation analysing anthocyanin vacuolar inclusions (AVIs) bodies in *5gt* mutant as a genetic background. *5gt* mutants cannot deliver soluble anthocyanins to the vacuole via ligandin-dependent importer pathway but use an autophagy-related vesicular pathway leading to excessive accumulation of insoluble AVIs that can be quantified inside the vacuole microscopically (Pourcel et al., 2010) (**Fig. 2B**). AVIs of SA-treated plants were less abundant, smaller and more circular than AVIs observed in control plants treated just with AIC. It supports our previous observations with absorbance measurements of anthocyanin content. Surprisingly, the abundance of AVIs does not decrease significantly between 200 µM SA and 400 µM SA, indicating a saturation effect of 200 µM SA. On the other hand, the area and circularity of AVIs are extensively affected by increased concentration of SA. AVIs in SA treated *5gt* plants tend to be smaller and more rounded than AVIs observed in control AIC-only conditions. Results from experiments using treatment with exogenous SA are in accordance with our data from mutants with high endogenous SA (**Fig. 1**).

Our observations differ from what has been described in other plant species after treatment with exogenous SA, where SA had a positive effect on anthocyanin production, as mentioned above in this chapter. Additionally, Liu et al. (2020) showed that anthocyanin accumulation caused by infection with *Penicillium corylophilum* correlates with increased SA content in *Arabidopsis*. Additionally, they demonstrated that treatment with exogenous SA induces anthocyanin production, and that this effect is dependent on NPR1 (Liu et al., 2020). Thus, it appears that SA has a dual role in regulating anthocyanin accumulation. The effect of SA and its downstream signaling depends on environmental conditions. If plants are grown under AICs, high SA levels may inhibit anthocyanin accumulation. Conversely, if the plant is not under apparent stress, elevated SA may trigger a stress response that leads to anthocyanin accumulation.

We speculate that the observed correlation between low anthocyanin accumulation and high SA levels could result from an ongoing spontaneous hypersensitive response (HR) in SA-overaccumulating mutants. This is a common phenomenon in such mutants, including autophagy mutants, and may also involve potential metabolic feedback.

### 3.3 SA-triggered reduction of anthocyanin vacuolar accumulation occurs independently from anthocyanin vesicular transport to the vacuole

Both *pi4kβ1/pi4kβ2* and *exo70B1* mutants exhibit defects in endomembrane vesicular trafficking (Kang et al., 2011; Kulich et al., 2013). However, expression of *NahG* in the *pi4kβ1/pi4kβ2* background fully alleviates the inhibitory effect on anthocyanin accumulation by downregulating SA content. While SA hyperaccumulation is reduced by the *tn2* mutation in the *exo70B1* background (Zhao et al., 2015), this only partially relieves the anthocyanin accumulation defect (**Fig. 1**). It suggests a potentially more prominent role for EXO70B1 in overall anthocyanin accumulation. Based on Kulich et al. (2013), it is probable NPR1-independent mechanism for SA accumulation in this mutant (Kulich et al., 2013).

To verify whether vesicle-dependent trafficking of anthocyanins into the vacuole is affected by elevated SA levels in an EXO70B1-dependent manner, we used a collection of *5gt, 5gt/exo70B1*, and *5gt/tn2/exo70B1* mutants. As highlighted above, in the *5gt* mutant background, the major carrier-dependent pathway for anthocyanin delivery to the vacuole is disrupted, and transport relies on the vesicular/autophagy machinery. Indeed, AVIs in *exo70B1/5gt* adult plants, which have high SA content, were significantly less abundant than in the *5gt* single mutant, but normalized in the *exo70B1/tn2/5gt* triple mutant (**Fig. 3**).

Thus, SA downregulation in the *exo70B1/tn2* double mutant background restores the ability to accumulate anthocyanins, suggesting that SA acts upstream to inhibit anthocyanin accumulation independently of EXO70B1-mediated vesicular uptake into the vacuole. Overall, our results imply that high endogenous SA, characteristic of *exo70B1* and several other mutants used in this study, has an inhibitory effect on overall anthocyanin synthesis and accumulation, including the 5GT/ligandin-dependent carrier import pathway. These observations indicate that SA accumulation correlates with reduced anthocyanin levels through currently unknown mechanisms that affect the overall availability or synthesis of anthocyanins, independent of their vacuolar delivery pathways.

## 4 CONCLUSION

We showed that high endogenous SA (and possibly its downstream NPR1-independent signaling, as suggested by (Kulich et al., 2013)) competes with or inhibits anthocyanin accumulation independently of both the ligandin-dependent and vesicular import pathways to the vacuole. These observations contribute to our understanding of how activated pattern-triggered immunity may inhibit anthocyanin production.

## Data availability

Raw data will be made available on request.

## Acknowledgements

We thank Petra Fialová for her excellent support. Also, to Martin Potocký, Jana Št’ovíčková for their help and assistance during the research. We thank to the imaging facility at Institute of Experimental Botany AS CR v.v.i. Part of the work was carried out with the support of the Growth Facility (BC Core Facilities; IPMB BC CAS). We would like to acknowledge Grammarly®, which we used for the improvement of clarity, flow, and grammatical accuracy.

## Author contribution

MD – AVI bodies, SA treatment, SD/LD anthocyanin measurement, analysing of the data, design and writing of the manuscript; OI – anthocyanin measurement, figures preparation; JSOH – anthocyanin and salicylic acid measurement; PBT – anthocyanin measurement, VM – anthocyanin measurement, preparation of *5gt/exo70B1* and *5gt/exo70B1/tn2* mutants; VZ – design of experiments; editing of the manuscript; TP – design of experiments, writing of the manuscript; MJ – design of the experiments, analysing of the data, writing of the manuscript. All the authors commented on the manuscript before its finalizing and approved the final version.

